# Allo: Accurate allocation of multi-mapped reads enables regulatory element analysis at repeats

**DOI:** 10.1101/2023.09.12.556916

**Authors:** Alexis Morrissey, Jeffrey Shi, Daniela Q. James, Shaun Mahony

**Affiliations:** Center for Eukaryotic Gene Regulation, Department of Biochemistry and Molecular Biology, The Pennsylvania State University, University Park, PA, USA

## Abstract

Transposable elements (TEs) and other repetitive regions have been shown to contain gene regulatory elements, including transcription factor binding sites. Unfortunately, regulatory elements harbored by repeats have proven difficult to characterize using short-read sequencing assays such as ChIP-seq or ATAC-seq. Most regulatory genomics analysis pipelines discard “multi-mapped” reads that align equally well to multiple genomic locations. Since multi-mapped reads arise predominantly from repeats, current analysis pipelines fail to detect a substantial portion of regulatory events that occur in repetitive regions. To address this shortcoming, we developed Allo, a new approach to allocate multi-mapped reads in an efficient, accurate, and user-friendly manner. Allo combines probabilistic mapping of multi-mapped reads with a convolutional neural network that recognizes the read distribution features of potential peaks, offering enhanced accuracy in multi-mapping read assignment. Allo also provides read-level output in the form of a corrected alignment file, making it compatible with existing regulatory genomics analysis pipelines and downstream peak-finders. In a demonstration application on CTCF ChIP-seq data, we show that Allo results in the discovery of thousands of new CTCF peaks. Many of these peaks contain the expected cognate motif and/or serve as TAD boundaries. We additionally apply Allo to a diverse collection of ENCODE ChIP-seq datasets, resulting in multiple previously unidentified interactions between transcription factors and repetitive element families. Finally, we show that Allo may be particularly effective in identifying ChIP-seq peaks in younger TEs, which hold evolutionary significance due to their emergence during human evolution from primates.

## Introduction

Next generation sequencing technologies underlie the study of various regulatory genomic phenomena, including gene expression (RNA-seq), protein-DNA interactions (ChIP-seq), 3D chromosome organization (Hi-C), and many more. While longer read sequencing techniques like Oxford Nanopore and PacBio have been established, many regulatory genomics assays continue to use short-read sequencing technologies due to the higher sampling rate (i.e., higher numbers of reads) and constraints with various preparation steps prior to sequencing. For example, the immunoprecipitation step in ChIP-seq is unlikely to allow the pull down of long stretches of chromatin and thus produces DNA fragments that are most compatible with short-read sequencing. Unfortunately, repetitive regions pose problems for short-read alignment. Any sequence that is repeated in the genome and is longer than the sequencing read length will create multi-mapped reads (MMRs). Using the common read lengths (35-75bp) seen in many studies and public repositories, 12-30% of sequenced reads are not uniquely mappable^1^ (Figure 1A). MMRs, due to their ambiguous nature, are generally removed during pre-processing in most regulatory genomics pipelines including those used by the ENCODE consortium^2^. For these reasons, repetitive regions have been largely overlooked in most gene regulatory analyses.

**Figure 1:**
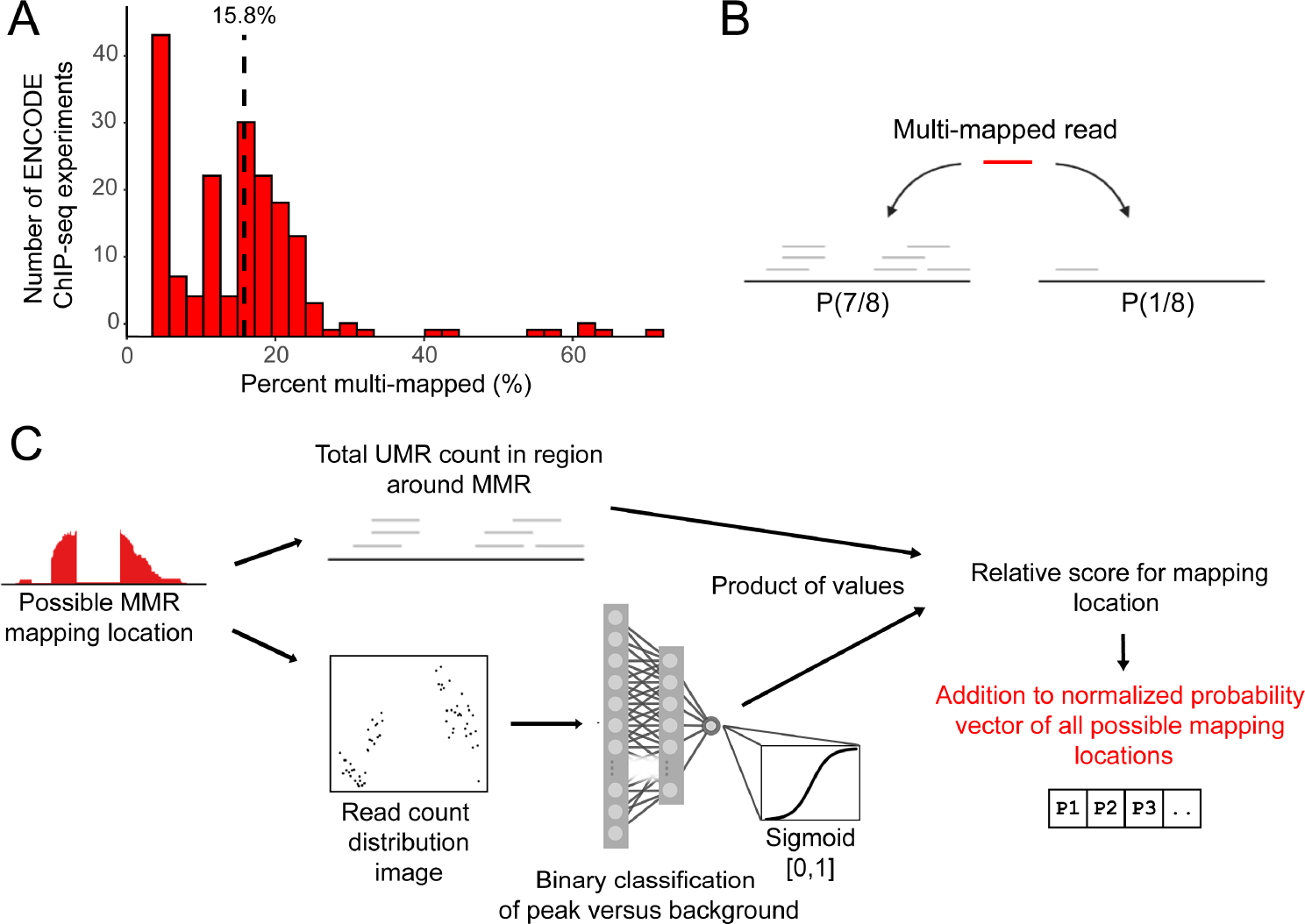
**A)** Proportions of reads that are multi-mapped across 200 K562 ENCODE datasets. **B)** Overview of probabilistic multi-mapped read allocation by uniquely mapped read count. **C)** Overview of the Allo algorithm combining uniquely mapped read counts and image classification of read distribution.

Several studies have demonstrated that repetitive regions contain transcription factor binding sites, suggesting that they play active roles in gene regulation^3,4^. One study of 26 orthologous transcription factors in mouse and human showed that around 20% of all binding sites were derived from transposable elements (TEs)^3^. Additionally, transposable elements have been shown to play a role in evolutionary adaptation. A prominent example is the insertion of a transposable element within the *cortex* gene of peppered moths^5^. This mutation increased transcription of the *cortex* gene which allowed for darker wing colors and better camouflage during the industrial revolution. TEs have also been implicated in the evolution of mammalian pregnancy development^6^, interferon response^7^, and pluripotency maintenance^8^. Transposable element mobilization has also been shown to perturb gene regulation and create disease states such as breast cancer^9^. Interestingly, most of these studies did not consider multi-mapped reads in their analyses and thus only investigated repetitive elements that were uniquely mappable. Therefore, the representation of transposable elements and other repetitive elements in regulatory processes is likely under-characterized.

Various methods have been proposed to deal with multi-mapped reads in gene regulatory data. One of the first methods mapped ChIP-seq reads directly to the consensus sequence of transposable elements and other repetitive regions^10^. While consensus sequence mapping provides family-level associations, these approaches cannot identify regulatory events at individual repetitive elements. Another set of approaches were developed to specifically allocate MMRs in the genome by using uniquely mapped read (UMR) counts in the vicinity of possible MMR mapping locations^11–13^. The intuition underlying these approaches is that a region containing UMRs is more likely to also have generated MMRs compared with alternate mapping locations. For example, MuMRescueLite^11^ used the UMR counts to create a simple probabilistic mapping of MMRs as illustrated in Figure 1B; MMR reads are allocated to specific locations based on the ratio of UMR counts at each of their possible mapping locations. Similarly, another method, CSEM, combined probabilistic mapping with iterative reweighting using expectation maximization^12^. Partial read counts are added based on this mapping in each iteration. Unfortunately, both CSEM and MuMRescueLite are, at time of writing, unusable due to uncompilable code and execution errors, respectively. The recent SmartMap method revived the probabilistic MMR mapping approach using a Bayesian model^13^. Due to the use of a Bayesian model over the entire genome, even fully mappable regions are modeled and thus UMR counts can be affected, introducing unnecessary error in the read coverage landscapes output by SmartMap. Additionally, SmartMap cannot be applied to single-end sequencing data without modifying the source code^13^.

Finally, all probabilistic MMR mapping methods to date have outputs that are not easily integrated into most regulatory genomics pipelines as they output custom file types^11^ or files types not accepted by commonly-used peak-callers^12,13^. In this work, we address the drawbacks of current approaches while also increasing the accuracy of MMR allocation. Our method, Allo, combines probabilistic mapping based on UMR counts with a convolutional neural network (CNN) that has been trained to identify the appearance of peak-containing regions. An overview of Allo’s method is shown in Figure 1C. Allo is usable for both single-end and paired-end sequencing data. Furthermore, the output of Allo is easily integrated into any pipeline as the output is an alignment file (SAM/BAM).

## Results

### Allo: multi-mapped read allocation combining probabilistic read allocation and image-based peak detection

Previous methods to allocate multi-mapped reads focused solely on UMR counts in regions around MMRs. Our method, Allo, also implements a probabilistic MMR mapping approach similar to that deployed by MuMRescueLite^11^ wherein regions with more UMRs will have a higher probability of being allocated an MMR. However, Allo combines probabilistic MMR mapping with a separate neural network module that predicts whether each possible MMR mapping location has read distributions consistent with a ChIP-seq peak (Figure 1B). The intuition behind this second module is that ChIP-seq reads are more likely to be generated from peak regions than from alternative non-peak locations on the genome.

Allo’s neural network takes the form of a convolutional neural network (CNN) that is trained on images of UMR distributions at peaks (Supplemental Figure 1). The CNN examines each possible mapping location of a given MMR and outputs scores ranging from 0 to 1, with higher values corresponding to CNN predictions that a region contains a UMR distribution consistent with ChIP-seq peaks. The total UMR count in each region is then multiplied by the corresponding CNN output score. Thus, areas with more UMRs and a more peak-like distribution of reads will have higher combined scores. Allo normalizes across the scores for each possible MMR mapping location to create a relative probability vector. The MMR is then allocated to a single possible mapping location by rolling a dice weighted according to the probability vector; i.e., locations with higher normalized scores are more likely to be allocated the MMR. A summary of Allo’s algorithm is shown in Figure 1C.

### Training a neural network to predict partially mappable peaks based on images of uniquely-mapped read distributions

ChIP-seq reads are distributed around protein-DNA binding events according to a characteristic shape, which is apparent at individual binding sites in successful ChIP-seq experiments given high enough sequencing coverage. Image-based classification approaches that leverage the shape properties have been previously used in multiple ChIP-seq peak calling applications^14,15^. We hypothesized that adding a measure of peak potential based on UMR distribution would increase the accuracy of MMR allocation within ChIP datasets. To train a neural network to predict peak potential, we obtained 10 transcription factor ChIP-seq datasets and 9 histone ChIP-seq datasets from ENCODE (Supplemental Table 1). The former were used to train a neural network on narrow peaks and the latter were used to train a neural network on mixed peaks.

To create a test set with known labels, we needed MMR-containing regions in which the peak status (i.e., peak versus not peak) is known. We achieved this by artificially shortening ENCODE ChIP-seq reads to create MMRs. The ground truth peak regions were defined as 500bp windows centered on MACS2 peak midpoints that were called from the full-length dataset. The 5’ read count distributions of artificially shortened UMRs in these peak windows were used to create 100x100 pixel images for CNN training. Treating the ChIP-seq data as images allows the CNN’s convolutional layer to find shape-related features at peaks without dependence on ChIP-seq signal levels. To make images for the negative set, we used an equivalent number of randomly selected MMR-containing regions from the background. A summary of this process is shown in Figure 2A. The binary classification accuracies using a cutoff value of 0.5 on the training datasets were 94.4% (narrow) and 92.3% (mixed), suggesting the neural networks learned to recognize partially mapped peaks. A summary of the test set generation process is shown in Figure 2A.

**Figure 2:**
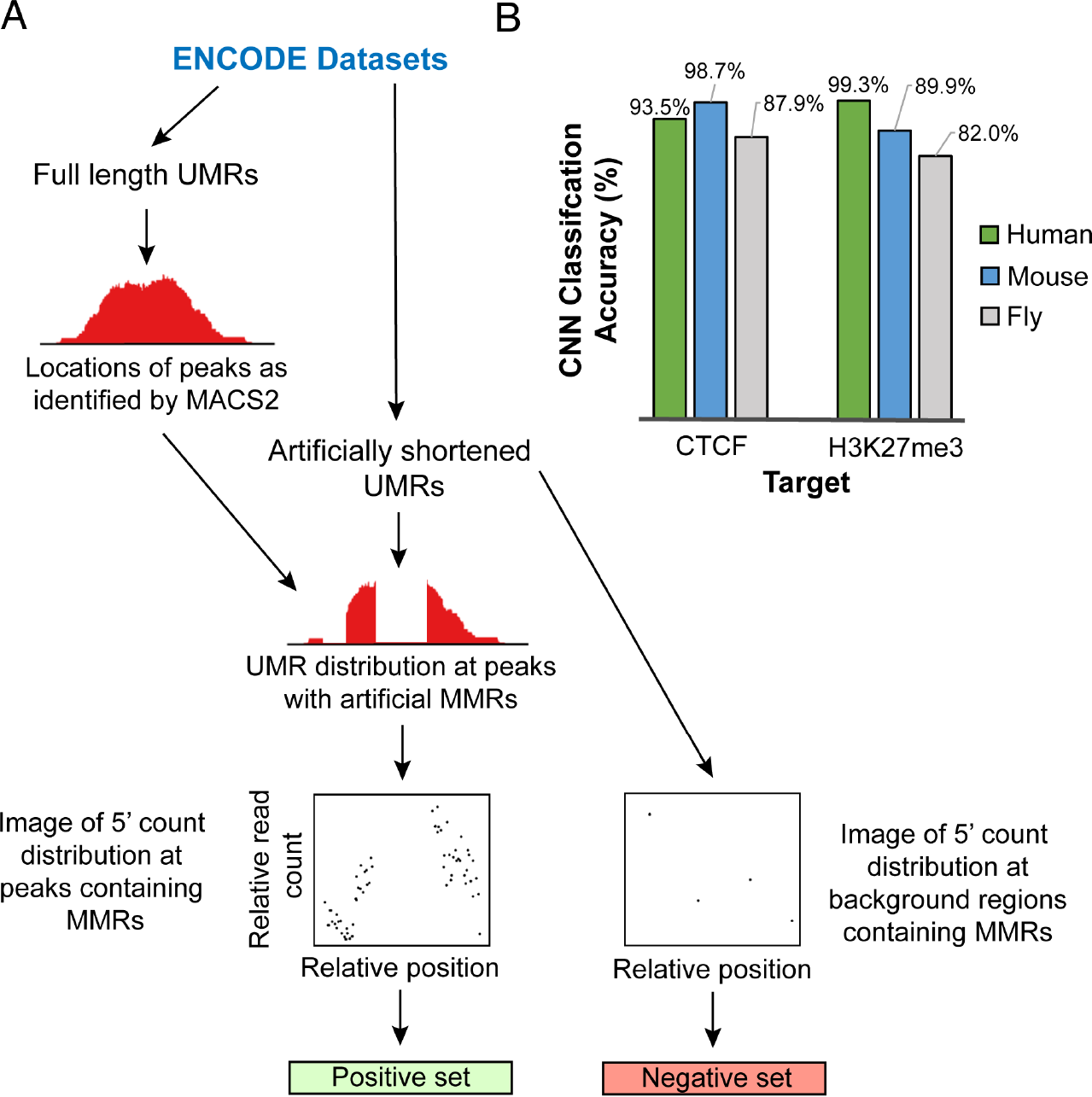
**A)** Overview of Allo’s neural network training process on ENCODE datasets. **B)** Binary classification accuracies of Allo’s neural networks on test datasets across species.

Our goal in designing Allo was to allow users to allocate multi-mapped reads easily across datasets. Therefore, we tested the ability of our neural networks to classify peaks in MMR-containing regions across ChIP-seq targets and species not used during training. These ChIP-seq datasets included CTCF and H3K27me3 across three species (human, mouse, and fly). Both narrow and mixed peak neural networks were able to predict above 80% accuracy (0.5 cutoff) across all datasets (Figure 2B) including those in other species.

### A CNN trained on read count distribution features increases accuracy of multi-mapped read allocation

To test the overall accuracy of Allo’s MMR allocation against other methods, we developed an approach using real ChIP-seq datasets while also having an estimate for ground truth. Briefly, we used a standard UMR pipeline to align reads and perform peak calling on a series of ENCODE ChIP-seq datasets. We then artificially shortened the reads (Figure 3B) and realigned them to the genome. Some of the reads that were uniquely mappable at full length then became multi-mapped at this new shortened length. For reads in this category, we had a ground truth location. Following alignment of the shortened reads, we used three separate methods to allocate MMRs: random allocation, read count only (similar to MuMRescueLite^11^), and Allo. We were unable to use CSEM^12^ as the software was no longer executable (see Methods for additional details). We were also unable to test SmartMap^13^ using this methodology as it outputs a BedGraph file which would not enable us to get accuracy on a per-read basis.

**Figure 3:**
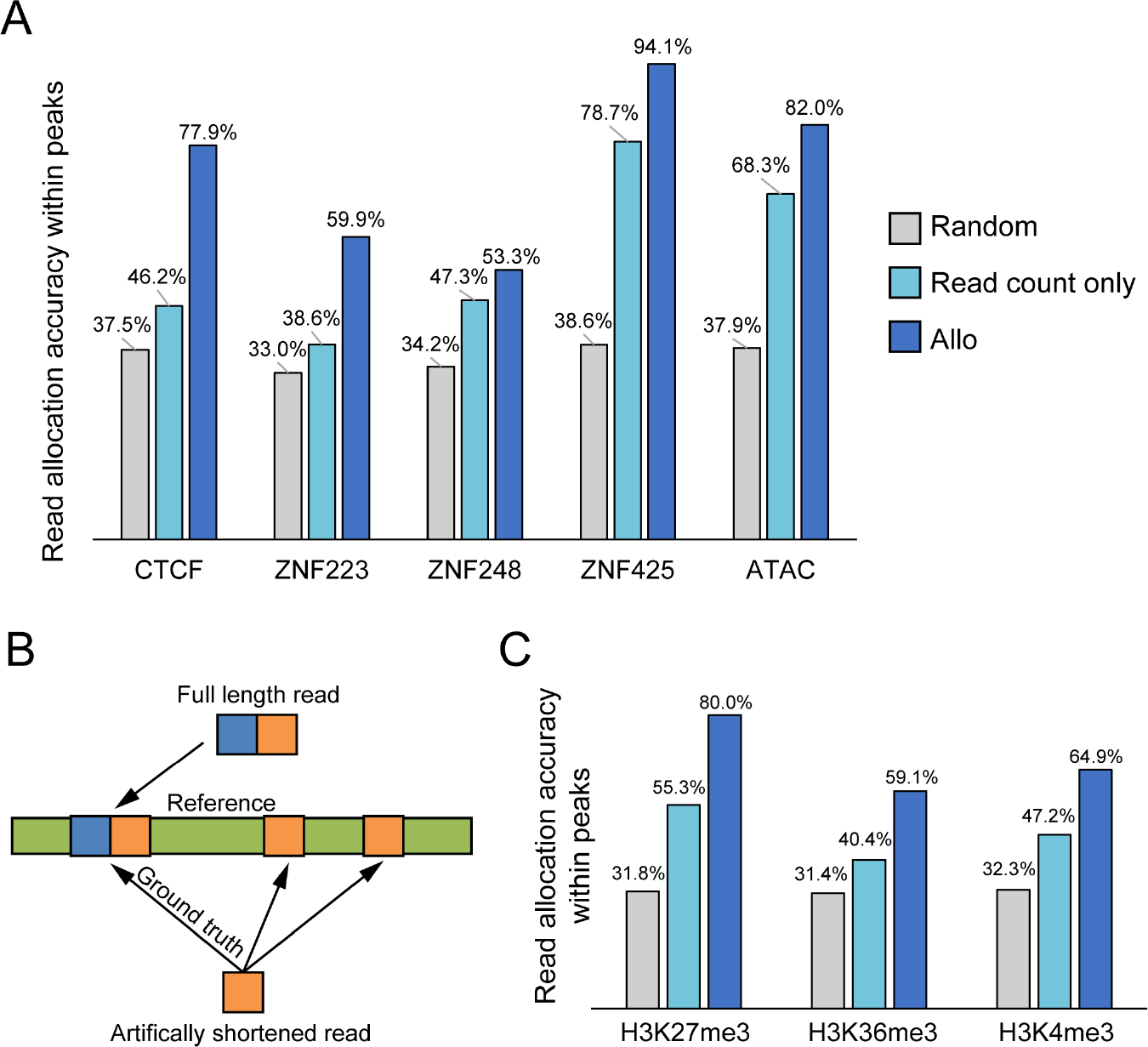
**A)** Accuracy of multi-mapped read assignment within peaks for selected K562 TF ChIP-seq datasets. Three different methods are tested: random assignment, read count only, and Allo in narrow peak mode. **B)** An overview of how the test datasets were generated by artificially shortening full length datasets. **C)** Accuracy of multi-mapped read assignment within peaks for selected K562 histone modification ChIP-seq datasets. Three different methods are tested: random assignment, read count only, and Allo in mixed peak mode.

Given the known interaction between CTCF and transposable elements^10^, we tested our methodology using a CTCF ChIP-seq dataset. In addition, we included three KRAB zinc-fingers of unknown function to emphasize the utility of Allo. KRAB zinc-fingers are associated with transposable element repression^4^ and thus we expected that these datasets would have peaks in repetitive regions. To ensure Allo is applicable across datasets, this set of transcription factors was excluded from the training set and we also chose datasets from different cell-types (Supplemental Table 2). As seen in Figure 3A, Allo outcompetes both random allocation as well as read count only allocation in all K562 datasets using narrow peak mode. To further show the broad applicability of our neural network across peak-based regulatory assays, we also show that Allo outcompetes other methods on an ATAC-seq dataset.

To test the mixed peak mode in Allo, we obtained ChIP-seq datasets for histone modifications H3K4me3, H3K27me3, and H3K36me3 (Figure 3C). Allo shows increased accuracy compared to the other two methods in these samples. Our results indicate that image-based classification using UMR distributions increases the accuracy of MMR allocation in both transcription factor datasets as well as histone modification datasets. Switching between narrow and mixed peak modes is a simple one argument option in the current implementation of Allo.

### Allo’s allocation accuracy and performance is competitive with that of SmartMap

As previously noted, we were unable to compare the read allocation accuracies of Allo and SmartMap, as it was not feasible to extract individual read mapping locations from the BedGraph files output by SmartMap. We were also unable to compare the two methods on a peak level basis, as MACS2 was unable to properly call peaks on SmartMap BedGraph files. Applying the MACS2 bdgbroadcall function to the SmartMap BedGraph files, as suggested in the SmartMap manuscript, resulted in over a million peaks called for each ENCODE dataset tested. Even if MACS2 had functioned properly, the bdgbroadcall does not allow for a control file and thus is not ideal for statistically valid analysis of ChIP-seq datasets. The SmartMap manuscript employed average read depths across simulated peaks to compare the absolute error of SmartMap and other methods. However, we sought to avoid using simulated ChIP-seq datasets in our comparisons, as it is difficult to accurately simulate realistic distributions of UMRs and MMRs around peaks.

To overcome these issues, we used our previous method of shortening full-length datasets and calculated the average read depth across peaks. We first removed all MMRs in the shortened datasets that were not UMRs in the full-length datasets, meaning all MMRs should have a ground truth location. We then ran SmartMap and Allo on these filtered alignment files. The BedTools map function was then used to calculate the average read depth at peaks, analogous to the comparison procedure performed in the SmartMap manuscript (see Methods).

As shown in Figure 4A, Allo had similar or lower percent error than SmartMap on all datasets tested. The only dataset in which SmartMap had a lower percent error than Allo was ZNF248 with a 3% advantage. The zinc-finger datasets have lower numbers of peaks than other datasets shown (Supplemental Table 3), and lower “fraction of reads in peak” (FRiP) scores (Figure 4A). It appears that in datasets with fewer peaks or lower FRiP scores, SmartMap and Allo tend to give similar results, whereas Allo is more accurate in datasets with higher signal-noise ratios (Figure 4A). This is not surprising given that Allo relies on the characteristic shape of UMRs at ChIP-seq peaks, and this shape will become more prominent in datasets with higher signal levels.

**Figure 4:**
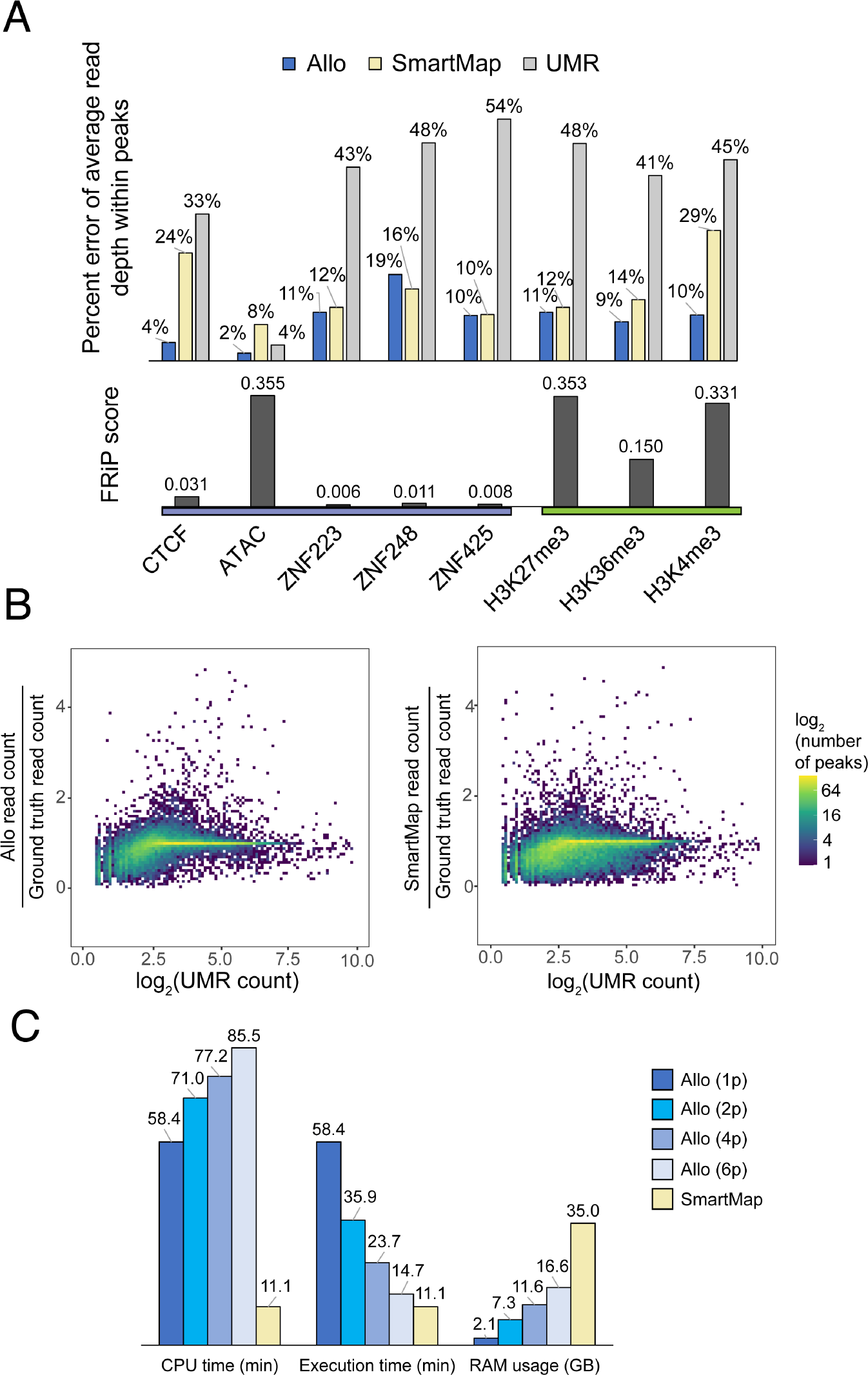
**A)** Percent error of the average read depth within peaks using Allo and SmartMap (top half) as well as the corresponding FRiP scores (fraction of reads in peaks). Datasets using the narrow peak mode of Allo are labeled as purple and the broad peak mode datasets are labeled as green. **B)** Percent error of Allo and SmartMap within peaks when compared to the log_2_(UMR) counts in those regions. **C)** Performance comparisons between Allo using various numbers of processes versus SmartMap. Values given are for datasets with 50 million reads.

Since both Allo and SmartMap rely on probabilistic assignment using UMR counts, we reasoned that regions containing fewer UMRs would be under-allocated multi-mapping reads. In Figure 4B, we show the percent error at each peak (combining all target samples used above) as a function of the total UMR count at each peak. As expected, regions with fewer UMRs are more likely to be under-allocated MMRs. However, Allo displays reduced error across all the datasets tested as seen by less discrepancy from the ground truth values overall.

Finally, we compared the computational requirements of the two methods using the test datasets described above. As shown in Figure 4C, Allo uses significantly more CPU time than SmartMap. This is likely because Allo is implemented in Python while SmartMap is implemented in C/C++. To mitigate the performance disparity, we implemented a multi-threading approach in Allo. Across the datasets tested above, Allo begins to approach the execution wall-time of SmartMap at around 6 processes. We also found that Allo uses significantly less memory than SmartMap. Due to unshared memory between processes, Allo will use more RAM when users choose a higher number of processes. Even so, Allo uses half as much memory as SmartMap on average at 6 processes. It is important to note that we ran SmartMap with default settings which includes only one algorithm iteration. We would expect that including more reweighting iterations would increase the execution time of SmartMap. Together, these results suggest that Allo is competitive with SmartMap in allocation accuracy as well as performance.

### Allo peaks have similar characteristics to uniquely-mapped peaks, reinforcing their validity

To demonstrate the utility of Allo and MMR-inclusive ChIP-seq pipelines, we focused on the properties of K562 CTCF peaks that are discoverable with the incorporation of Allo but are not detected using a traditional UMR-only ChIP-seq analysis pipeline. Allo’s inclusion of MMRs uncovered 3,114 CTCF peaks in addition to the 54,677 found using UMRs alone (Figure 5A). Peaks only found using Allo are labeled as “Allo-only” peaks. Allo increased the read depth significantly in these regions as seen in Figure 5B. Because CTCF binds strongly to its cognate motif, we also performed motif analysis on both sets of peaks. The CTCF motif was the highest ranked motif found by MEME-ChIP in both the UMR peaks (Figure 5D) and the Allo-only peaks (Figure 5E), suggesting that these peaks are characteristic of true CTCF binding sites. The cognate motif was also centrally enriched in the Allo only peaks as seen in Figure 5C using a heatmap of CTCF motif TRAP scores at each peak^16^. Due to CTCF’s presence at TAD boundaries, we examined the overlap between UMR peaks and Allo-only peaks at these boundaries using TAD calls from the 3D Genome Browser^17^. A TAD boundary was defined as being within 2.5kb of the beginning or end of a TAD. We found that there was a similar overlap percentage between peak sets and TAD boundaries; 22.4% for the UMR peaks and 19.1% for the Allo-only peaks. Figure 6A shows an example of two Allo-only CTCF peaks at TAD boundaries near the THOC3 gene. In this case, the MMRs present are a result of a duplicate pseudogene of THOC3. Figure 6 shows a higher resolution image of these peaks demonstrating that the allocation of MMRs enabled peak discovery. Allo peaks tend to have the same characteristic read distribution shape as peaks found in uniquely-mappable regions. Together, each of these results suggest that peaks found using Allo are authentic.

**Figure 5:**
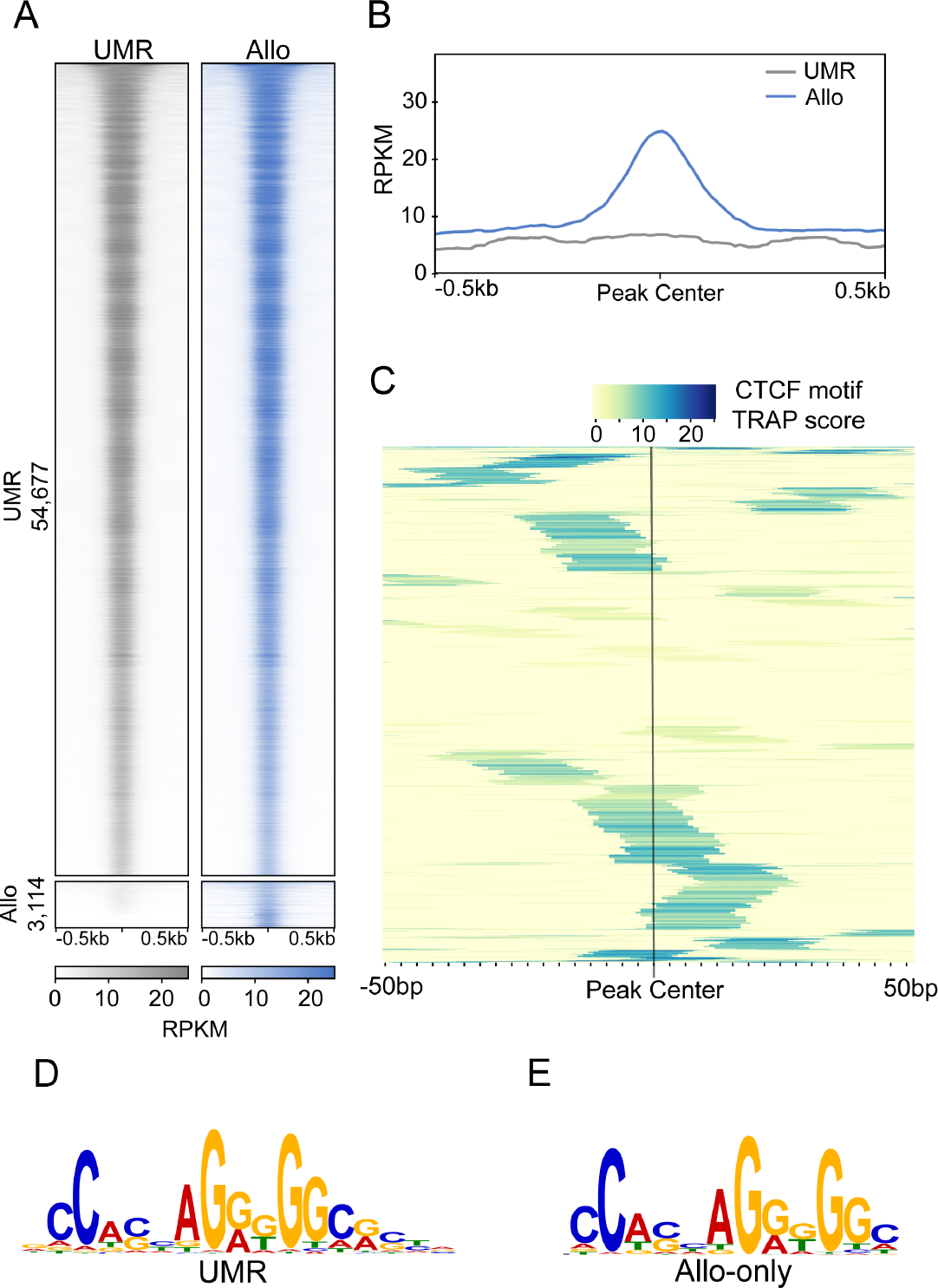
**A)** ChIP-seq heatmaps comparing peaks called by MACS2 using UMRs only or UMRs plus Allo-mapped reads. **B)** Read depth at Allo-only peaks using UMR BAM files versus with Allo output. **C)** TRAP scores of the CTCF motif within 3,114 peaks only found using Allo-mapped reads. Sorting of peaks is based on hierarchical clustering of rows. BAM files. **D)** and **E)** Top ranked motifs from MEME-ChIP in the UMR and Allo-only peaks, respectively.

**Figure 6:**
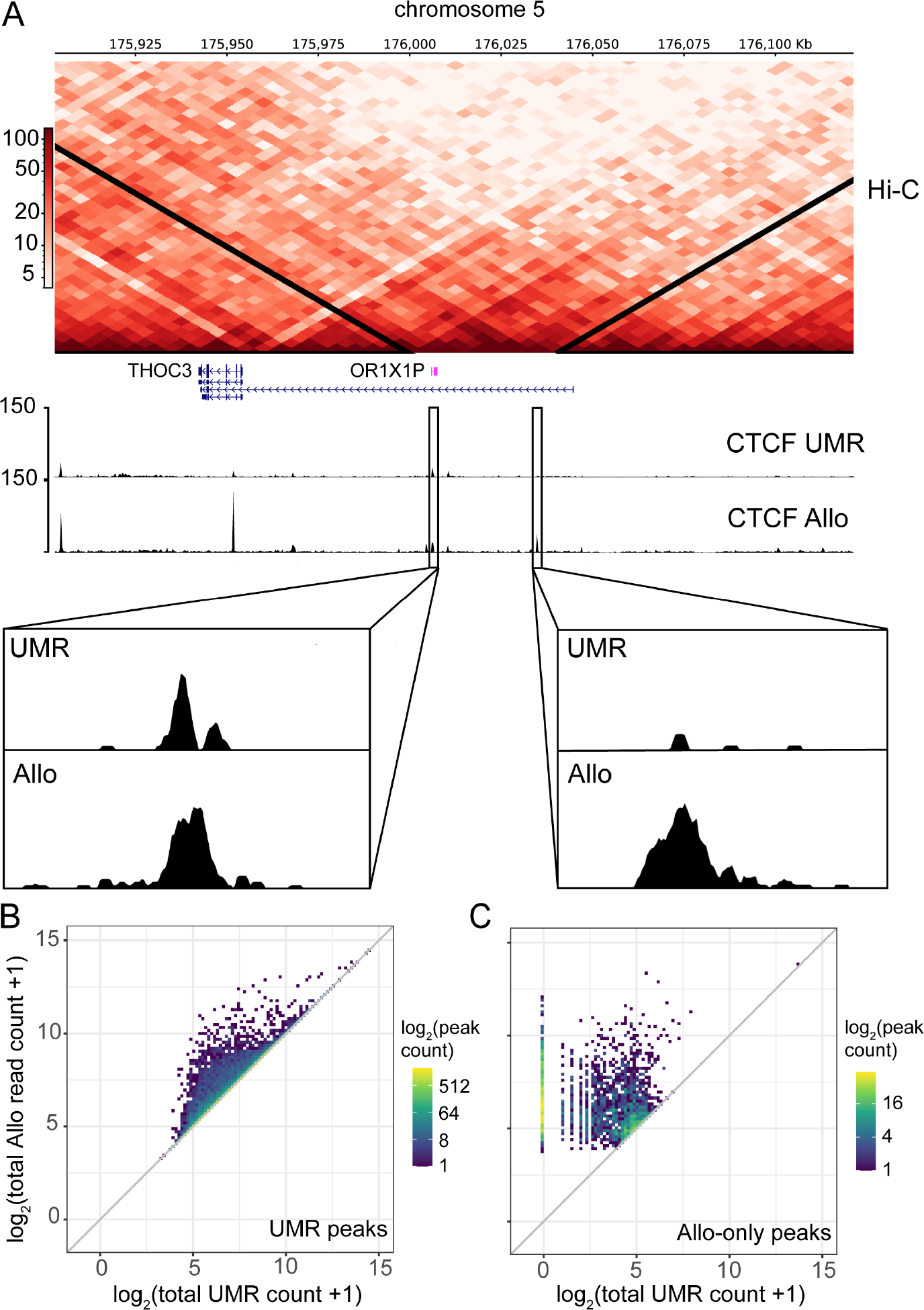
**A)** Hi-C interaction heatmap showing two TAD boundaries on chromosome 5 near the THOC3 gene. Read counts are also plotted for the UMR-only analysis and the Allo analysis. Bottom sections show the zoomed in version of two Allo-only peaks at these at these TAD boundaries. **B)** Density scatterplot showing the total read count before and after the incorporation of Allo within UMR peaks. **C)** Density scatterplot showing the total read count before and after the incorporation of Allo within Allo-only peaks. Note that that Allo was run using default settings which allocates MMRs even in cases where there are no nearby UMRs. This results in peaks called in regions with 0 starting UMR counts as seen in panel C. The user can shut off this functionality using the argument “--remove-zeros” to have more stringent criteria for MMR allocation

Furthermore, we investigated the effects of Allo on peaks that were discoverable using only UMRs, as some of these peaks additionally contain MMRs. Figures 6B and 6C show the total read counts before and after using Allo, at UMR peaks and Allo-only peaks, respectively. It is evident that many UMR peaks gain substantial numbers of reads with the inclusion of MMRs. Therefore, Allo can increase peak resolution and ChIP-seq enrichment quantification accuracy at many peaks, even those that were already discoverable using UMR-only pipelines. Supplemental Figure 3 provides examples of the increased resolution at selected UMR peaks after MMR inclusion.

### Allo supports the discovery of associations between TFs and repeat families in large scale ChIP-seq analyses

To broadly survey how an MMR-inclusive pipeline might enable the discovery of additional protein-DNA binding events, we re-analyzed 200 ENCODE K562 ChIP-seq datasets using Allo. The use of Allo resulted in a mean increase of 13.5% additional peaks compared with the standard UMR-only pipeline (Figure 7A). We next evaluated the repetitive element content of these newly discovered Allo-only peaks by overlapping with RepeatMasker^18^ annotated repetitive regions. Allo-only peaks have a higher overlap with repetitive elements compared with UMR peaks (Figure 7B, each dot represents a ChIP-seq dataset). A majority of the tested datasets have points above the diagonal, demonstrating that Allo-based analysis finds more peaks in repetitive elements than UMR pipelines alone. It is notable that there are samples in which Allo-only peaks have proportionally less overlap with repetitive elements. This can be explained by the various other causes of MMRs that would not be annotated as repetitive elements. For example, BDP1 and BRF1 were among the farthest from the diagonal (27.9% and 20.3% decrease, respectively, in repeat element overlap for Allo-only peaks). Both proteins are part of RNA Polymerase III transcription initiation factors. RNA polymerase III primarily transcribes tRNAs and thus the MMRs present in these analyses would not coincide with RepeatMasker-annotated repetitive elements, even though tRNAs appear in multiple similar copies across the genome.

**Figure 7:**
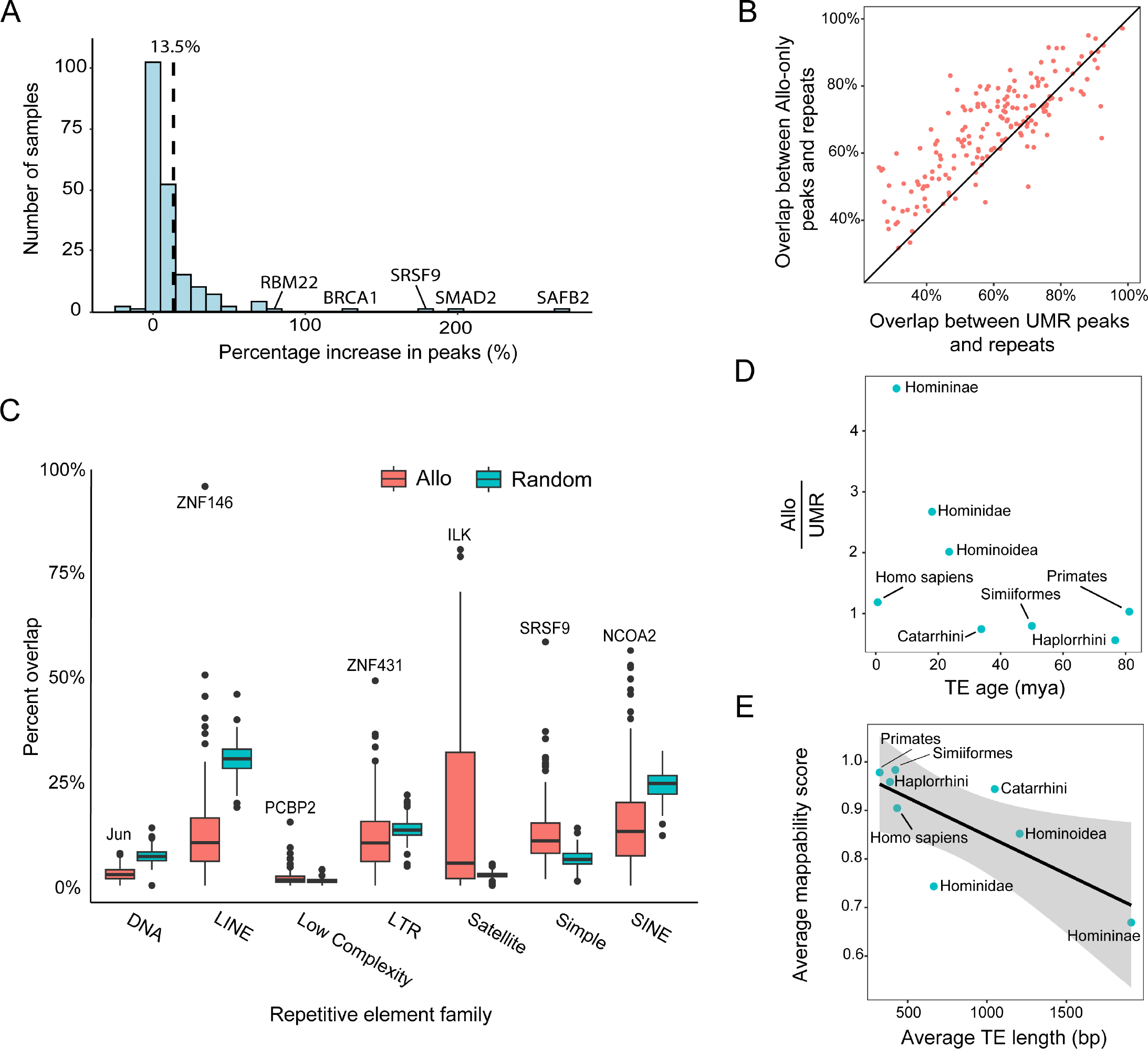
**A)** Percentage increase in peaks between the Allo-inclusive pipeline and the UMR-only pipeline across 200 ENCODE K562 ChIP-seq datasets. **B)** Percentage overlaps between Allo-only peaks and RepeatMasker-annotated repeats compared with the same comparison in UMR peaks. **C)** Percent overlap between Allo-only peaks and various repetitive element families. The plot also shows the overlap rates for random regions with comparable region lengths. The highest outliers are labeled in each set. **D)** Comparison of the increase in overlap of various transposable element families between UMR peaks and Allo-only peaks split according to transposable element age. Most recent common ancestors are labeled. **E)** The average mappability scores versus average transposable element length for each TE’s most recent common ancestor.

To further investigate Allo’s ability to find new peaks in repetitive elements, we analyzed the associations between Allo-only peaks and each of the major classes of repeats. For each analyzed ENCODE dataset, we compared repeat-class overlap rates between the Allo-only peaks and randomly sampled genomic regions of the same number and lengths (Figure 7C). From this analysis, we observed that many of the repetitive element families were underrepresented compared with their occurrence on the genome. This general underrepresentation is not necessarily surprising; TF binding sites may be lacking in the large set of repeats that are located in heterochromatin regions, or the underrepresentation could point to a cohort of binding sites in highly repetitive regions that remain undetectable even after using an MMR-inclusive analysis.

The exceptions to the overall underrepresentation were satellite repeats and simple repeats. Satellite repeats are commonly found within centromeric and telomeric regions^19^ and have been shown to regulate gene expression in eukaryotic organisms^20–22^. While these elements may thus contain abundant TF binding sites, it is also possible that the overrepresentation seen here is an artifact of assembly limitations. Due to their repetitive nature, many satellite repeats are not present in common assemblies such as hg38, which we used for our analyses. It is possible that the satellite repeat copies present in hg38 become sinks for multi-mapped reads that would have been assigned to background regions had all copies been present in the assembly. For future projects, the telomere to telomere genome assembly could be used to mitigate problems associated with repeat representation and further elucidate the relationships between all classes of repeats and gene regulatory elements^23^.

While most of the repetitive element families were underrepresented compared to random regions, there were outliers within the ENCODE datasets. Extracting these outliers could be used as an exploratory tool to identify transcription factors that bind to specific repetitive element families. For example, we found that the transcription factor Jun (part of the AP-1 complex) was highly bound to DNA repeat elements. A previous study found that DNA repeat elements like MER44B had an overrepresentation of motifs for various immune related factors including AP-1, which is activated by inflammatory cytokines.^24^ Other notable outliers were the BTB zinc-finger TFs ZNF146 and ZNF431, which overlapped strongly with LINE-1 and LTR repeats, respectively. Unlike the known association between KRAB zinc-fingers and LINE-1 elements, there is no prior evidence to suggest an interaction between these BTB factors and particular repeat elements. Both factors could be explored further to identify their functions and the role transposable element binding plays in those functions.

### Allo helps to identify peaks in younger transposable element families

Previous studies using UMR-only pipelines have nevertheless found significant levels of overlap between transcription factor binding sites and various transposable elements^3^. We hypothesized that while UMR pipelines can detect peaks in many transposable element families, Allo would be better suited to identify peaks in younger families as we can assume they are less diverged and thus less uniquely mappable. To test this hypothesis, we intersected all Allo-only peaks within our ENCODE datasets with groups of repeat families based on their most recent ancestor. We found that newer families generally tended to have more Allo-only peaks when compared to older families (Figure 7D), supporting the use of Allo for studying recent repeat-derived regulatory divergence. Interestingly, the general pattern was not followed by human-specific transposable elements, the most recent repetitive elements in the genome. We would assume the mappability of these newer human-specific elements would be lower than primate elements, but the mappability of these elements based on UMAP K100 values was in fact quite high (Figure 7E). One reason for this outlying observation could be the types and average lengths of elements that are human-specific. Longer transposable elements are less mappable as read lengths are less likely to extend past them. Human-specific elements, on average, appear to be much shorter than many of the other age groupings (Figure 7E), explaining the discrepancy we see within Figure 7D.

## Discussion

While repetitive elements have long been recognized to play a role in gene regulation^25^, there has not been a concerted effort to include multi-mapped reads in standard sequencing analyses. As a result, many regulatory elements within repeats remain invisible to regulatory genomics assays. A major reason for this exclusion is that existing methods to allocate multi-mapped reads are not easily integrated into established analysis pipelines. For instance, the recent SmartMap method produces a BedGraph file, making it incompatible with standard pipelines like those used by ENCODE^2^. Older methods, such as MuMRescueLite and CSEM have not been maintained and are no longer functional, and they also lacked read-level output. Allo addresses these shortcomings by producing an alignment file, ensuring compatibility with various downstream analyses.

Besides increasing basic usability, we also wanted to create a method that was competitive with previous approaches. Random allocation is used frequently in pipelines as it is the easiest method to implement; both BWA and Bowtie perform random allocation by default. Allo significantly outcompetes random allocation as seen in Figure 3. Allo also has greater read assignment accuracy compared with a strategy based entirely on UMR-weighted probabilistic mapping of MMRs (similar to the method implemented in MuMRescueLite; Figure 3), or the Bayesian mapping approach implemented in SmartMap (Figure 4A). These comparative results demonstrate the advantages of Allo’s integrated neural network peak classifier, which aims to take advantage of UMR distribution shapes in alternate mapping locations.

To emphasize the importance of including multi-mapped reads in ChIP-seq analyses, we analyzed 200 ENCODE datasets using Allo. The average total peak number increase was 13.5% (Figure 7A), resulting in thousands of additional peaks identified in many of the samples. Five datasets in this sample actually doubled their peak numbers after the inclusion of Allo including BRCA1, a well-known tumor suppressor gene^26^. In analyzing repetitive element family associations (Figure 7C), we also identified other TFs involved in cancer progression, including Jun^27^, ILK^28^, SRSF9^29^, and NCOA2^30^, that are highly associated with particular repeat families. Outside of Jun, none of these TFs have been previously identified as binding to repetitive elements. We additionally found that Allo was especially important for the discovery of binding sites in younger transposable elements (Figure 7). Previous studies have identified primate-specific enhancers that may have contributed to the evolutionary development of humans^31,32^. Enhancers that arose from repetitive elements are less likely to be categorized in these types of studies due to issues with multi-mapped read inclusion. Thus, Allo could be used to further study the regulatory contributions of transposable elements in an evolutionary context. Together, the results of this work suggest that rescuing multi-mapped reads with Allo should be used across all datasets, even those without hypotheses tied to repetitive regions. The inclusion of MMRs in future analyses may finally elucidate the function of these overlooked areas of the genome.

## Methods

### Availability

Allo is available under an open source license (MIT license) from https://github.com/seqcode/allo. Allo can also be installed from PyPI using “pip install bio-allo” at the command line. Allo is also available as a bioconda package: https://anaconda.org/bioconda/allo. Instructions on usage can be found in the above GitHub repository.

### Datasets

All ChIP-seq and ATAC-seq datasets were obtained from ENCODE^2^ with the exception of the *Drosophila* H3K27me3 ChIP-seq dataset, which was obtained from NCBI GEO. Supplemental Table 1 lists the IDs for datasets used in training Allo’s CNN. Supplemental Table 2 lists the IDs of all datasets used for testing. Supplemental Table 3 lists the IDs of all K562 datasets extracted from ENCODE in our large exploratory analysis.

### Percentage of multi-mapped reads

To find the percentage of multi-mapped reads in ENCODE datasets, we used the single-end FASTQ files listed in Supplemental Table 3. All FASTQs were aligned using Bowtie2 v.2.5.0 and percentage multi-mapped was extracted from the output of Bowtie2 upon alignment.

### Convolutional neural network training and testing set generation

To train the neural network, we first obtained 10 transcription factor ChIP-seq datasets and 9 histone ChIP-seq datasets (Sup. Table 1). These datasets were aligned to hg38 using Bowtie1 v.1.0.0 with the arguments “--best -- strata -m 1 -k 1 --chunkmbs 1024”. This represented our full length read set and MACS2 v.2.2.7.1 was used to call peaks with all default parameters. These were our ground truth peaks. We next artificially shortened these reads from the 3’ end using Cutadapt v.4.2 with the parameters “-u -LENGTH”. The lengths trimmed from the reads varied dependent on the original read lengths and are shown in Supplemental Table 1. We used Bowtie v.1 instead of Bowtie v.2 due to Bowtie v.1s’ ability to align very short reads more accurately after this artificial trimming. Next, we aligned the artificially shortened reads to hg38. For the UMR samples, we used the following arguments “--best --strata -m 1 -k 1 --chunkmbs 1024”. For the MMR samples, the arguments were “--best --strata -m 25 -k 25 --chunkmbs 1024”. We then used Allo’s parser utility (argument is --parser) to separate out the UMRs and MMRs in the shortened samples.

To find the 5’ read counts within these peaks and background regions, we created a custom script (cnn_training_gen.py in the Allo GitHub repository). The user must supply the script with four arguments: the alignment (SAM) file; a bed file of peak regions; a bed file of background regions; and the output name. To make the positive and negative bed files, we first took the MMR only artificially shortened SAM file (output of Allo’s parser utility) and converted it into a bed file. We then overlapped this bed file using Bedtools v.2.27.1^33^ intersect -u with the ground truth peak file. This gave us the positive regions. For the negative regions, we used Bedtools intersect -v to find areas with MMRs that did not overlap ground truth peak regions. These regions have different lengths but this is corrected to +/-250bp from the midpoint within the Python script. Additionally, the Python script also corrects the negative set by only outputting an equal amount to the positive set. The output is a text file with comma separated arrays of 5’ read counts in 500bp windows around the positive and negative regions. The labels are also output with a tab separator in between. The bash script for this entire pipeline can be found in the Allo GitHub repository (cnn_training_examples.sh). The same script was used to generate the testing set.

### Convolutional neural network architecture and training

The two neural networks used in Allo’s allocation algorithm are image based. To convert the 5’ read count text files explained in the above section into images, we used a custom Python function. This can be found on the Allo GitHub repository within the first code block of the Python markdown file (Narrow_train.ipynb). We trained both neural networks using Tensorflow v.2.11.0. Both neural networks were trained using a batch size of 500 and had the same architecture. The basic architecture of the neural networks can be found in Supplemental Figure 1. The optimizer used was Adam and the loss function used was binary crossentropy. The specific code used to train the neural networks can be found on the Allo GitHub repository under files named Narrow_train.ipynb and Mixed_train.ipynb.

### Allo allocation algorithm

Allo’s algorithm has two main phases. The first phase is the pileup of uniquely-mapped reads within the genome. To do this, Allo first loops through the alignment file and parses uniquely and multi-mapped reads. Depending on the aligner used, alignment files can contain locations that do not have the highest alignment score and thus require extra parsing. In addition, Allo identifies the correct pairs when using paired-end sequencing data. When Allo encounters a uniquely-mapped read, it adds it to a Python dictionary. The keys are the locations on the genome and the values are the 5’ read counts in that specific location. A dictionary was used in order to increase the speed of data acquisition to create images of various regions on the genome in phase II. For paired-end reads, only the first read in each pair is used to construct this dictionary. The second in pair can be used instead through the argument “--r2”. Also, during this phase, temporary files are constructed from multi-mapped reads.

In phase II, the multi-mapped reads in the temporary files are allocated. Allo analyzes one read at a time by grouping it with its possible locations. Two vectors are constructed. The first vector contains the total read count in 500bp windows (+/- 250bp) around each location. In this vector, 1 is added to all location UMR counts so that locations with 0 UMRs are still considered. The second vector contains the output of the sigmoid function from Allo’s neural network. The 5’ read counts in 500bp windows around each read location are converted to an image using the dictionary described above and a custom script to bin and plot the counts. This script can be found on the Allo GitHub repository within the file Narrow_train.ipynb. After constructing the image, it is fed through the neural network and the results are stored in the vector above. The result is two vectors with each index corresponding to a location that the read being analyzed has mapped to. These vectors are then multiplied together to get a final score vector. The final score vector is then normalized by dividing all entries but the sum of the vector giving the final probabilities. The choice function in Python is then used to select the location based on the final probabilities.

To increase the speed of Allo’s algorithm, a few extra steps were employed. Firstly, Allo stores previously analyzed regions in a dictionary as the reanalysis slowed down Allo considerably. In addition, Allo does not create an image for regions with 0 read counts as the result will be consistent across these regions. Rather, it stores the value for 0 count regions once. These regions are stored as NULL in the dictionary above. Finally, Allo does not create an image for regions containing fewer than 3 reads. The reasoning behind this was similar as described for 0 regions. When we created 1 million randomized regions with 3 reads in a 500bp window, the results were very similar from the neural network across the many images. We believe this is because low read depth regions do not have the resolution that allows Allo’s neural network to make useful predictions. The average across 1 million random 3 count total regions was 0.0062 from the final sigmoid function. This average value is thus pre-defined as the neural network output for all regions with 3 or fewer total read counts.

### Read allocation accuracy

To measure the allocation accuracy of various methods, we needed to create a peak set that contained multi-mapped reads but had a ground truth read count. FASTQ files were downloaded from ENCODE for 7 experiments (Supplemental Table 2). To avoid problems with mixing single and paired-end data, experiments that were paired-end were treated as single-end by only utilizing the first read file (this is only for the purposes of accuracy testing; Allo can properly handle paired-end data as described above). All files were aligned using Bowtie v. 1.0.0 with the parameters “--best --strata -m 1 -k 1 --chunkmbs 1024”. Bowtie v.1.0.0 was used because of its higher accuracy in aligning very short reads. The reads were then artificially shortened from the 3’ end using Cutadapt (4.2) with the parameters “-u -LENGTH”. The lengths trimmed varied based on original length and can be found in Supplemental Table 2. The shortened reads were then aligned with Bowtie v. 1.0.0 using the same parameters as above. The full-length alignments were used to call peaks using MACS2 v.2.2.7.1 on all default parameters. This became the ground truth peak set. Using Bedtools v.2.27.1 intersect, we extracted the reads in the full-length dataset that fell within the ground truth peaks. This gave us our final ground truth alignment file needed for comparisons.

Allo (default), Allo (--read-count) and Allo (--random) were then used to allocate the reads in the artificially shortened samples. The locations of these allocations were compared to the alignment file from the full-length dataset. The accuracy percentage was simply calculated as the total number of correctly allocated reads divided by the total number of MMRs in the sample. The full script for this calculation including an example from ENCODE is available on the Allo GitHub (allo/scripts/read_acc.sh).

### SmartMap comparison

#### Read depth percent error

FASTQ files were downloaded from ENCODE (IDs listed in Supplemental Table 2). Paired-end reads were aligned with Bowtie2 v.2.5.0 to hg38 using the arguments “-k 25 --no-mixed --no-discordant.” UMRs were extracted from the resulting alignment files using the parser utility in Allo (--parser). These UMRs were then used to get ground truth peak locations using MACS2 v.2.2.7.1 with the “-f BAMPE” argument. We next artificially shortened these reads from the 3’ end using Cutadapt v.4.2 with the parameters “-u -LENGTH”. The lengths trimmed from the reads varied depending on the original lengths and are shown in Supplemental Table 1. The use of Bowtie2 in this section was due to its ability to more accurately map paired-end reads even with the pitfall of it being less accurate for short read lengths. We then used the bash join function to extract MMRs from the shortened sample that had a ground truth in the full-length sample. Many of the MMRs in the shortened samples were also MMRs in our ground truth full-length sample and we wanted to avoid these in our final calculations. The resulting alignment files were then analyzed by both SmartMap and Allo. To use an alignment file with SmartMap’s prep script, we had to make some modifications. The modified version of this script can be found on the Allo GitHub under the name sm_prep.sh. To convert the Allo output to a bedgraph file, we used Bedtools v.2.27.1 bamtobed -bedpe and then bedtools genomecov -bga. We used this same conversion on the ground truth full-length alignment file.

To get the read depth within peaks, we used bedtools map with the arguments “-c 4 -o mean -null “0”“. The bed file used was the peaks identified by MACS2 for the full-length datasets. The percent error was then calculated for each peak and the average was taken for each sample. An example script of this pipeline is available on the Allo GitHub named smartmap_compare.sh.

#### FRiP score calculation

To calculate the FRiP scores, Bedtools v.2.27.1 intersect -u was used to find the overlap between peaks called by MACS2 in the sample and the associated alignment files. Samtools v.1.16.1 view -c was then used to get the number of reads within this overlap. This number of reads was then divided by the total number of reads to calculate the FRiP score for each specific sample.

#### Performance metrics

CPU usage, execution time, and RAM usage were all tested on Intel Xeon Gold 6226R CPUs with a processing speed of 2.90GHz. We ran Allo using various numbers of processes (1,2,4, and 6) as well as SmartMap using the 7 testing datasets from the above section. We normalized the values to get the metrics at 50 million reads total, making the performance metrics comparable across datasets. The mean was calculated from all 7 datasets and is what is shown in Figure 5C.

### CTCF analysis

#### Alignment and peak calling

FASTQ files for CTCF in K562 were downloaded from ENCODE. Samples files were ENCFF000YLW and ENCFF000YLY. The control file used was ENCFF000YRB. Reads were aligned using Bowtie v. 1.0.0 to hg38 with the arguments “--best --strata -m 25 -k 25 --chunkmbs 1024.” Reads were allocated using Allo on default parameters. We extracted the UMRs using grep against the ZA and ZZ tags from Allo. We then called peaks on the UMR alignment file as well as the Allo alignment files using MACS2 (2.2.7.1) with default parameters. Peaks were called on the replicates separately and overlapping peaks between the replicates found using Bedtools v.2.27.1 intersect -u were used as the peak sets. Using Bedtools intersect -v we were able to identify peaks only discovered via Allo.

#### Heatmap and profile plot

Both the heatmap and the profile plot were created using DeepTools v.3.5.1^34^. The bamCoverage function was used to create bigwigs from each of the alignment files. We used the RPKM normalization argument. To check that this normalization was valid for our specific datasets, we plotted profile plots of random regions using Bedtools random (Supplemental Figure 2). We used the computeMatrix function with the arguments “--referencePoint center -a 1000 -b 1000” before plotting.

#### Motif identification and scanning

Using Bedtools v.2.27.1 getFasta with hg38, we extracted sequences for all peaks represented in the UMR sample as well as Allo-only peaks. We then input these sequences into MEME-ChIP v.5.3.3 with the arguments “-meme-nmotifs 5 -minw 5 -maxw 20”. The motifs in Figure 6 were those with the lowest p-values. To scan for the CTCF motif in the Allo-only peaks, we used the online version of TRAP v.1.0^16^. The matrix file given was JASPAR vertebrates and the background model was human promoters. The top result for TRAP motifs found was CTCF, as anticipated. We used a custom script to create 100bp arrays with the TRAP scores. The TRAP output table was used to get which region of the 100bp window for each peak contained the score. We then plotted these arrays using Seaborn v.0.12.2^35^ clustermap. Hierarchical clustering was performed by Seaborn only on the rows (peaks).

#### TAD overlap

The locations of TADS were downloaded from the 3D genome browser for K562, specifically the dataset from Rao et al. (2014)^17^. We then used this to create a bed file of TAD boundaries which we considered as within 50kb of either end of each TAD. This file was then intersected with UMR peak calls and Allo-only peak calls using Bedtools intersect -u. Figures were constructed using pyGenomeTracks^36^.

#### UMR and MMR count comparison

Bedtools v.2.27.1 multicov was used to calculate read counts at Allo-only peaks and UMR peaks. The BAM files used were concatenated files from both CTCF biological replicates. The data was then plotted using ggplot2 with geom_bin2d and a continuous scale with a log_2_ transformation.

### ENCODE K562 analysis

#### Alignment and peak calling

We analyzed the ENCODE public database, focusing specifically on ChIP-seq experiments in K562 cells. From 774 qualifying experiments (retrieval date Feb. 2023), we randomly selected a subset of 200 transcription factors for more in-depth analysis. We retrieved FASTQ files for both single- and paired-end experiments. Technical replicates were concatenated and aligned using Bowtie 2 v.2.5.1, reporting 25 valid alignments per read with parameters --no-mixed --no-discordant for paired-end reads. The alignments were subsequently collated using samtools v.1.16.1. We employed Allo on the sorted alignments to obtain rescued reads. For control experiments, reads were randomly assigned during Allo processing. Following alignment and rescue, peaks were subsequently identified using MACS2 v.2.2.7.1, with the BAMPE format designated for paired-end sequencing data. ENCODE blacklist regions were excluded using Bedtools v.2.27.1 intersect (-v). A list of all ENCODE dataset used can be found in Supplemental Table 4.

#### Overlap with transposable elements families

The RepeatMasker annotation file was used to get locations of repetitive elements families in hg38. Bedtools v.2.27.1 intersect (-u) was then used to find the overlaps between each ENCODE dataset and each repetitive element family. Bedtools random was used to construct random bed files that matched each ENCODE dataset. These randomized bed files had the same number of peaks with the same lengths as their matched ENCODE dataset. The overlaps between these bed files and repetitive element families was also found using Bedtools intersect (-u). The fraction overlaps were then plotted as a boxplot using ggplot2.

#### Transposable element dating

We used data from Simonti *et al*.^37^, which identified the most recent common ancestor for each transposable element family. Using this data, we created bed files that contained the repeats dating to each common ancestor. Each file contained the transposable elements and their locations (as identified by RepeatMasker annotation). Bedtools v.2.27.1 intersect (-u) was used to find the fraction overlap between Allo only peaks and UMR peaks for each common ancestor.

The data from Simonti *et al*. did not include information about years since divergence. To date the speciation events, we leveraged data from two separate studies. The first, from Pozzi *et al*.^38^, identified the years since speciation for Homo sapiens, catarrhini, haplorrhini, and primates. This was done using comparisons between mitochondrial genomes. The second study, from Perelman *et al*.^39^, identified speciation events using polymorphisms in specific genomic regions of interest. They identified the speciation time for homininae, hominidae, hominoidea, and simiiformes. For both studies, we took the highest bound for divergence time estimates. The speciation time for each common ancestor was then plotted against the ratio of overlap between Allo only regions and UMR only regions.

#### Transposable element mappability scores and lengths

A BedGraph file containing UMAP100^40^ scores for hg38 was downloaded from the Hoffman laboratory website found at “https://bismap.hoffmanlab.org/” To get the average score across each transposable element family, bedtools map was used (-o mean) against the RepeatMasker annotation file. Transposable elements from each common ancestor were extracted from this file and the average was taken across all transposable elements in each set. The average length of transposable elements was found by extracting locations for each common ancestor and taking the average of those location lengths within the bed file.

## Acknowledgements

This manuscript is based upon work supported by the National Science Foundation DBI CAREER 2045500 (to S.M.). Any opinions, findings and conclusions or recommendations expressed in this material are those of the authors and do not necessarily reflect the views of the National Science Foundation. The Mahony lab is also supported by the National Institutes of Health grant R35GM144135. A.M. gratefully acknowledges funding and training opportunities from the Computation, Bioinformatics, and Statistics (CBIOS) Training Program (T32GM102057). The authors thank the members of the Center for Eukaryotic Gene Regulation at Penn State for helpful feedback and discussions.

