## Supplemental Figures for "Allo: Accurate allocation of multi-mapped reads enables regulatory element analysis at repeats"

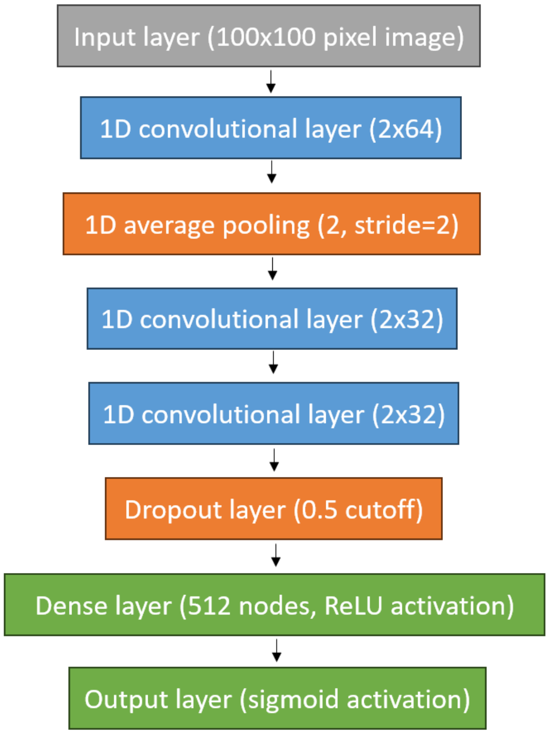


**Supplemental Figure 1:** Architecture of Allo's convolutional neural network. The structure is the same for both the narrow and broad peak CNNs.

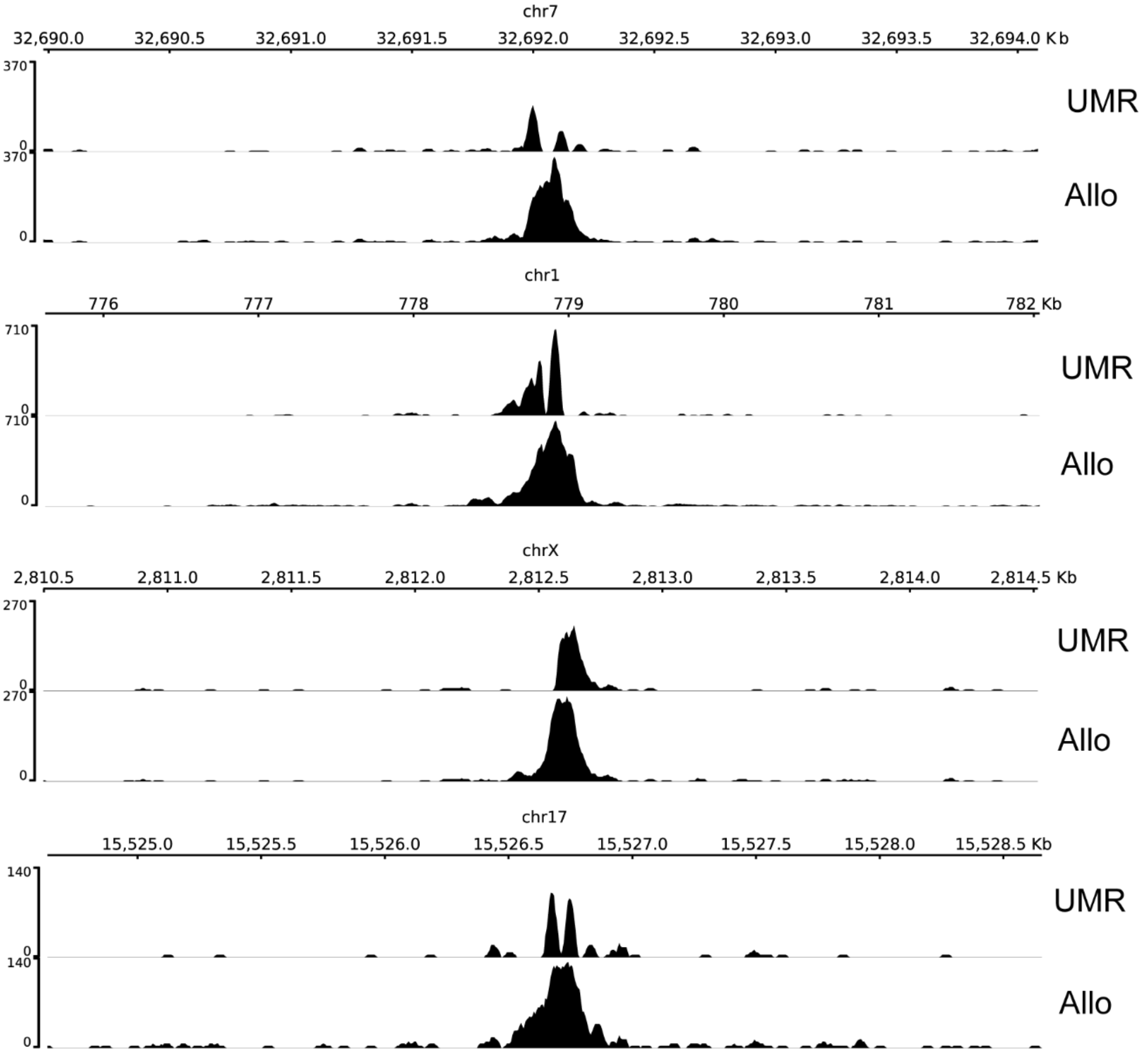


**Supplemental Figure 2:** Read pileups at locations of uniquely-mapped peaks that increased in resolution after the inclusion of multi-mapped reads. Pileups are labeled as either UMR for uniquely-mapped reads only or Allo for after the inclusion of multi-mapped reads.

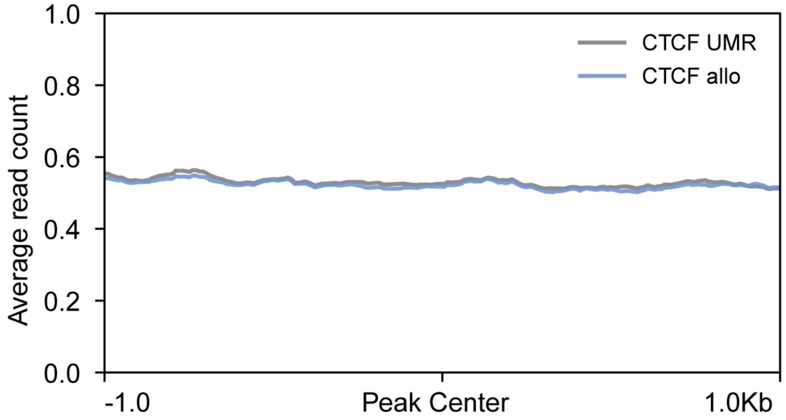


**Supplemental Figure 3:** Profile plot of read counts within random background regions in CTCF K562 dataset. Read counts are shown using UMR counts only as well as after the inclusion of Allo. RPKM was used for normalization by DeepTools for both datasets.
